# Clonal hematopoiesis driven by *Dnmt3a* mutations promotes metabolic disease development

**DOI:** 10.1101/2025.05.02.651715

**Authors:** Bowen Yan, Qingchen Yuan, Prabhjot Kaur, Annalisse R. Mckee, Daniil E. Shabashvili, Olga A. Guryanova

## Abstract

Clonal hematopoiesis (CH) is associated with an increased risk of non-hematologic chronic diseases including metabolic disorders, yet the causality remains poorly defined. *DNMT3A* is the most frequently altered gene in CH, commonly through monoallelic loss-of-function (LOF) and Arg882His (RH) hotspot mutations. Here we demonstrate in a mouse model that CH driven by *Dnmt3a* RH and especially LOF promotes obesity, diabetes, and chronic liver disease, further exacerbated by high-fat diet (HFD).

## Main

Clonal hematopoiesis (CH) is defined as clonal expansion of hematopoietic stem cells (HSCs) marked with somatic mutations and of their progeny in the absence of quantitative blood abnormalities. In addition to an elevated risk of developing blood malignancies, CH is notably associated with various chronic diseases beyond the hematopoietic system, such as inflammatory disorders and cardiovascular disease^1-5^. Observations from large epidemiological studies including UK Biobank found presence of CH was enriched in individuals with an increased waist-to-hip ratio, type 2 diabetes (T2D), and chronic liver disease^4,6,7^, although the degree of these effects varied between cohorts. Importantly, whether CH is a cause or a consequence of these comorbidities—a question of high translational significance—is incompletely understood.

*DNA methyltransferase 3A* (*DNMT3A*) is the most commonly mutated gene in CH (up to 40% of all cases)^8^. While the *DNMT3A* Arg882His (RH) hotspot variant is enriched in acute myeloid leukemia (AML), most *DNMT3A* alterations observed in CH are consistent with the loss-of-function (LOF)^9,10^, indicating these types of mutations may not be fully mechanistically and functionally equivalent. However, the specific impact of CH driven by different types of *DNMT3A* mutations on chronic disease development has not been investigated.

To elucidate the functional impact of CH with *DNMT3A* LOF and RH mutations in the hematopoietic system on metabolic disease development, we created a mouse bone marrow (BM) transplantation (BMT)-based chimeric model with 20% of either *Dnmt3a*^*+/-*^ (representing LOF), *Dnmt3a*^*+/R878H*^ (corresponding to human Arg882His), or *WT* control cells (marked by panleukocytic CD45.2) mixed with 80% *WT* support (CD45.1) to mimic CH clone^11-13^ (Fig. 1A, SFig. 1A-B). Eight weeks post-transplantation to allow hematopoietic engraftment and reconstitution, mice were randomized to receive high-fat high-glucose “western” diet (HFD) to induce metabolic disease or remained on normal chow as control. In the normal chow control condition, we observed that animals harboring a proportion of *Dnmt3a*^*+/-*^ or *Dnmt3a*^*+/RH*^ hematopoietic cells exhibited a significant increase of body weight gain over time and a trend toward higher food intake compared to *Dnmt3a*^*WT*^-engrafted controls (SFig. 1C-E, Fig. 1B). This was accompanied by subcutaneous white adipocyte hypertrophy (Fig. 1C-D). These effects were more pronounced in the *Dnmt3a*^*+/-*^ group compared to the *Dnmt3a*^*+/RH*^ chimerae. Further, HFD lead to more rapid bodyweight increase and greater extent of adipocyte hypertrophy, particularly in the *Dnmt3a*^*+/-*^ group (SFig. 1E, Fig. 1B-D). The *Dnmt3a*^*+/-*^ HFD group became overweight at 2 weeks and obese at 4 weeks on HFD, which is one week earlier than the *Dnmt3a*^*WT*^ HFD group (SFig. 1F). Compared to *Dnmt3a*^*WT*^ controls, *Dnmt3a*^*+/-*^-CH animals exhibited significantly higher plasma leptin and resistin levels after 6 weeks of HFD consistent with increased adiposity, whereas *Dnmt3a*^*+/RH*^-CH animals on HFD showed an increase in plasma resistin levels (Fig. 1E, SFig. 1G). Notably, 6 months after transplantation, corresponding to 4 months after randomization to remain on normal chow, both *Dnmt3a*^*+/-*^ and *Dnmt3a*^*+/RH*^-CH animals developed mildly elevated plasma leptin levels even without a dietary challenge (SFig. 1H-I), suggesting a direct obesogenic effect of *Dnmt3a*-CH. In line with a well-established link between overweight, insulin resistance, and diabetes, when maintained on HFD these mice exhibited a greater propensity to develop impaired glucose metabolism characterized by elevated fasting blood glucose levels, increased plasma insulin, and impaired glucose tolerance (Fig. 1F-H). Although when *Dnmt3a*^*+/-*^-CH mice were maintained on normal chow, their plasma insulin and glucose tolerance remained unperturbed, almost half of these animals had their fasting glucose levels fall within pre-diabetic range 7 months post-transplantation unlike other groups (SFig. 1J-L). These findings suggest *Dnmt3a*-CH, particularly driven by *Dnmt3a* LOF, directly contributes to metabolic dysfunction similar to obesity and T2D.

**Figure 1.**
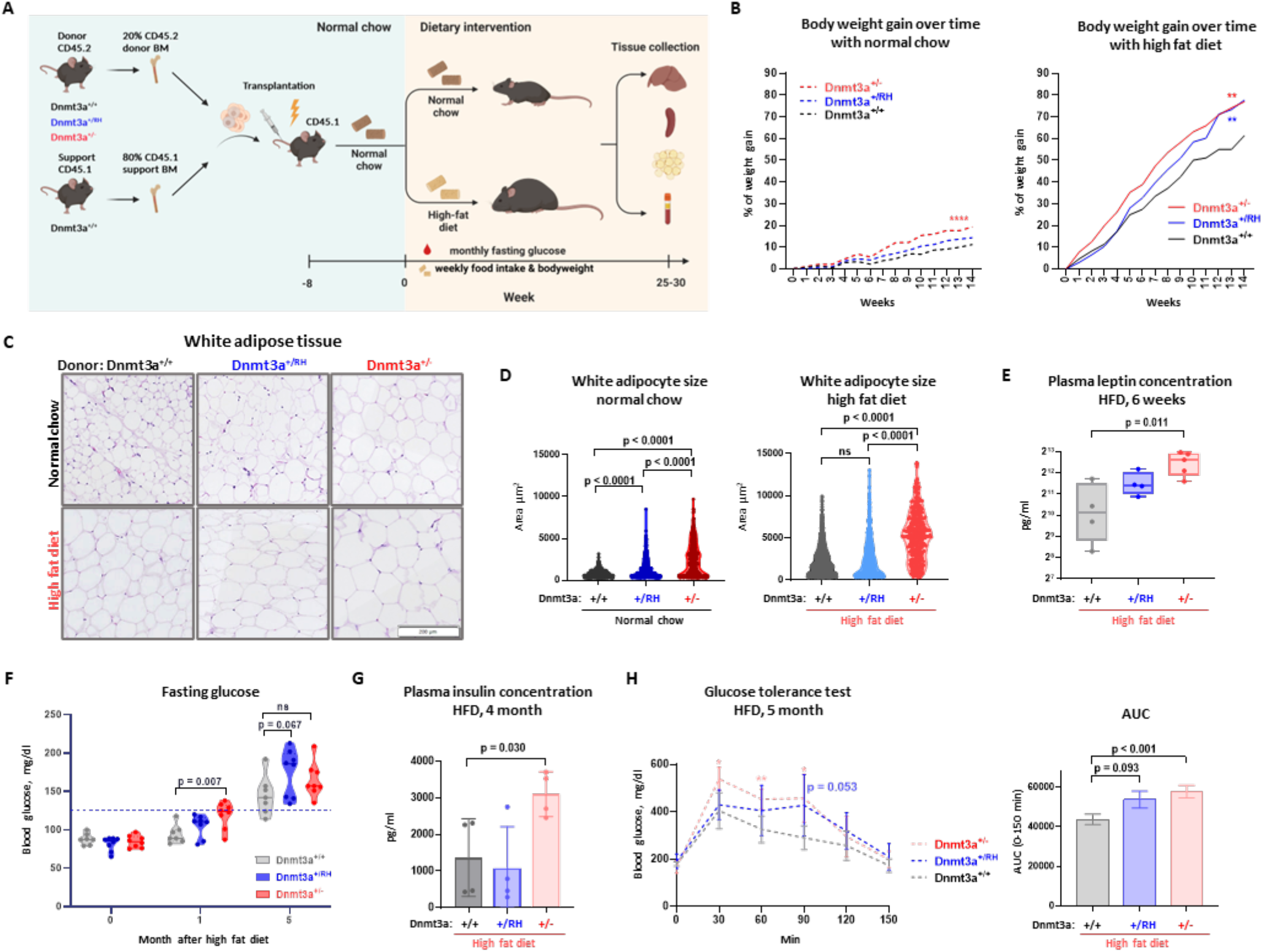
Hematopoietic-specific *Dnmt3a* alterations in a model of CH promote obesity and impaired glucose tolerance. (A) Experiment design (schematic created with BioRender.) (B) Animal body weight gain; mixed-effects analysis, **** *p*<0.0001, ** *p*<0.01 compared to control *Dnmt3a*^*+/+*^ group; n≥7 for all groups. (C-D) Hematoxylin and eosin (H&E) staining of white adipose tissue (C, bar – 200 µm) and adipocyte size (n=3 animals per group, 2 fields per animal). (E) Plasma leptin levels after 6 weeks of HFD. (F) Fasting glucose levels at baseline and after 1 and 5 months on HFD. (G) Plasma insulin concentration after 4 months on HFD. (H) Glucose tolerance test (GTT): blood glucose levels after glucose load in fasted animals and calculated area under the curve (AUC). Data are presented as mean ± standard error, n≥5 for all groups; * *p*<0.05, ** *p*<0.01 in GTT with Student’s *t*-test compared to *Dnmt3a*^*+/+*^ controls at individual time points.

Given the emerging link between CH, inflammation, and changes in myelopoiesis^14,15^, next we examined lineage composition in the hematopoietic system and cytokine profiles. *Dnmt3a*^*+/-*^-CH mice showed increased abundance of donor-derived inflammatory monocytes (CD45.2) in both spleens and peripheral blood compared to wild-type competitor cells (CD45.1) within the same recipient and to *Dnmt3a*^*WT*^-CH controls, most pronounced under HFD conditions; *Dnmt3a*^*+/RH*^-CH group exhibited intermediate phenotype (Fig. 2A, SFig. 2A). Notably, this phenotype was present in both *Dnmt3a*^*+/-*^ (CD45.2) and wild-type (CD45.1) compartment, reflecting a generalized pro-inflammatory state (SFig. 2B). Consistently, after 10 weeks of HFD, *Dnmt3a*^*+/-*^-CH mice had elevated levels of plasma inflammation-related cytokines such as IL-15 and MIP-1α, compared to *Dnmt3a*^*WT*^-engrafted controls under HFD conditions (Fig. 2B) although levels of classical pro-inflammatory cytokines such as IL-1β and IL-6 were less perturbed.

**Figure 2.**
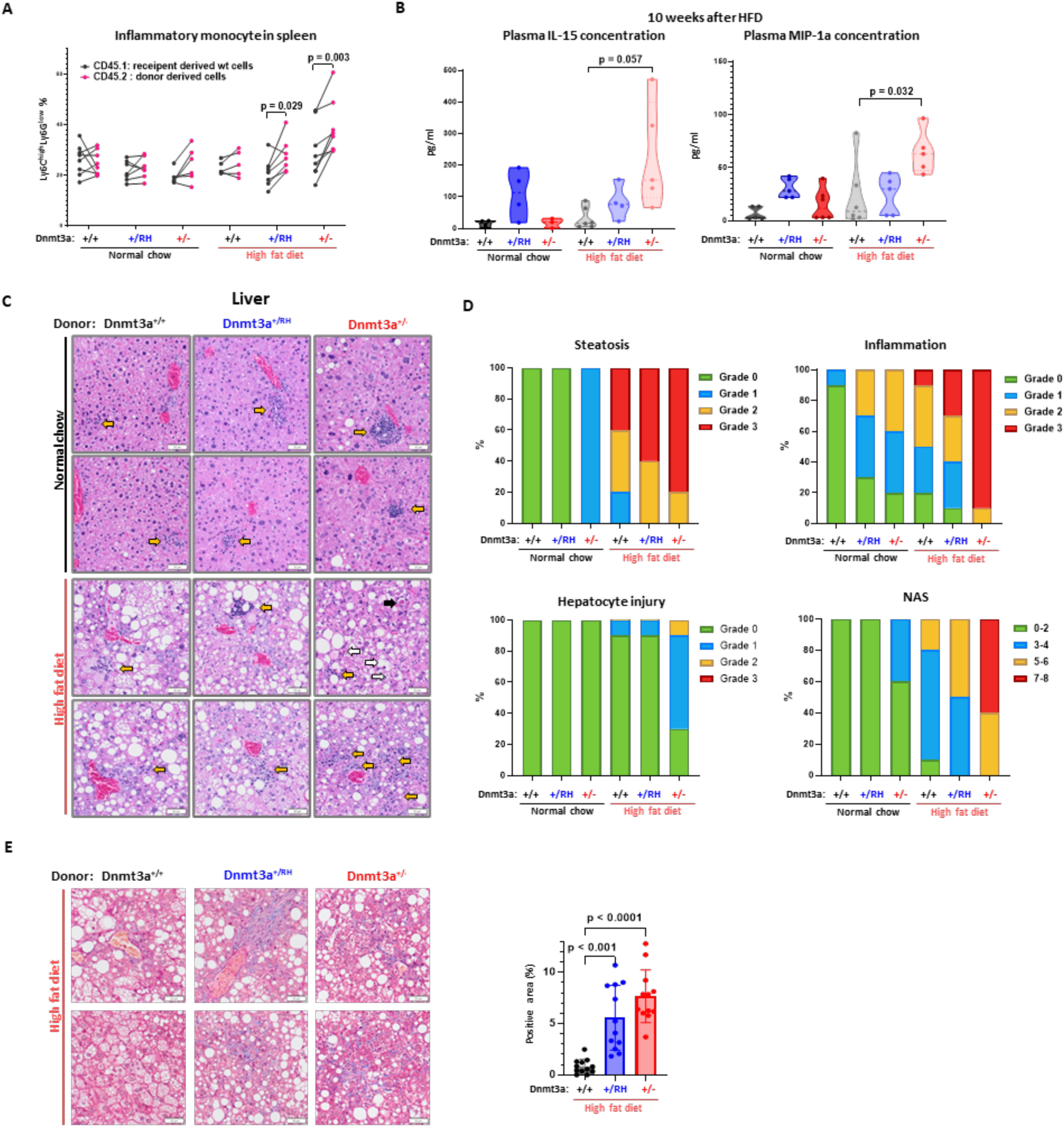
Model CH with *Dnmt3a* deficiency or mutation promotes inflammation, advanced steatohepatitis and liver damage. (A) Inflammatory monocytes in donor (CD45.2) or wild-type competitor/host (CD45.1) myeloid cells in spleen; paired Student’s *t*-test. (B) IL-15 and MIP-1α in blood plasma, unpaired *t*-test with Welch’s correction as appropriate. (C) Histopathological analysis of H&E-stained liver sections (black arrow: acidophilic body, white arrows: ballooned hepatocytes, yellow arrows: inflammatory infiltrates; bar – 50 µm.) (D) Steatohepatitis was graded using nonalcoholic steatohepatitis histologic criteria (n=3 per group, 5 fields per animal). NAS: Nonalcoholic Fatty Liver Disease Activity Score. (E) Fibrosis detected by Masson’s trichrome staining of liver sections and its extent quantified as percentage of blue-stained positive area; unpaired *t*-test with Welch’s correction (n=3 per group, 4 fields per animal).

The meta-inflammatory state was further evident in the livers of both *Dnmt3a*^*+/-*^-CH and *Dnmt3a*^*+/RH*^-CH animals. When maintained on normal chow, both *Dnmt3a*-CH groups exhibited greater lobular hepatic inflammation with prominent infiltrating immune cell aggregates compared to *Dnmt3a*^*WT*^ controls (Fig. 2C-D, SFig. 2C). Further, *Dnmt3a*^*+/-*^-CH animals developed signs of metabolic dysfunction-associated steatotic liver disease (MASLD), characterized by an increase in number and size of macrovesicular fat droplets (Fig. 2C-D, SFig. 2D). Under HFD, *Dnmt3a*^*+/-*^-CH group progressed to severe MASLD with marked hepatitis reflected by high non-alcoholic fatty liver disease activity score (NAS)^4,16^ (Supplementary Table 1) and extensive fibrosis (Fig. 2C-E).

Collectively, our findings indicate that in a mouse model, presence of CH driven by *Dnmt3a* LOF or RH promotes the development of obesity, impaired glucose metabolism similar to T2D, and chronic inflammatory liver disease reminiscent of MASLD in humans. The metabolic disease is further exacerbated by HFD, most strongly in the *Dnmt3a*^*+/-*^ context. Our study highlights the differential impact of various CH mutations and urges detailed mutation- and gene-specific investigation of CH in human chronic disease pathogenesis, risk stratification, and mitigation through pharmacologic, lifestyle, and dietary interventions.

## Contributions

B.Y. designed the study, performed experiments, analyzed data, and wrote the manuscript. Q.Y. performed experiments, contributed to data interpretation, and edited the manuscript. P.K., A.R.M. and D.E.S. performed experiments. O.A.G. conceived the study, supervised the study, co-wrote the manuscript, and secured funding. All authors revised and approved the final manuscript.

## Acknowledgements

This work was supported in part by NIH award R01DK121831 to OAG. OAG is also supported by the Edward P. Evans Foundation, the Oxnard Family Foundation. BY is supported by the Ocala Royal Dames for Cancer Research, Inc. and the ACS Institutional Research Grant to the University of Florida Health Cancer Center (UFHCC). UFHCC is an NCI-designated cancer center (P30CA247796). Flow cytometry analyses were performed at the UF Interdisciplinary Center for Biotechnology Research (ICBR) RRID:SCR_019119.

## Ethics declarations

All authors declare no competing financial interests.

**Supplementary material** consisting of Methods, Supplementary Figures 1 and 2, and Supplementary Tables 1 and 2 is provided as a separate document.

## References

1 Weeks, L. D. & Ebert, B. L. Causes and consequences of clonal hematopoiesis. Blood 142, 2235–2246, doi:10.1182/blood.2023022222 (2023).

2 Kessler, M. D. et al. Common and rare variant associations with clonal haematopoiesis phenotypes. Nature 612, 301–309, doi:10.1038/s41586-022-05448-9 (2022).

3 Jaiswal, S. et al. Clonal Hematopoiesis and Risk of Atherosclerotic Cardiovascular Disease. N Engl J Med 377, 111–121, doi:10.1056/NEJMoa1701719 (2017).

4 Wong, W. J. et al. Clonal haematopoiesis and risk of chronic liver disease. Nature 616, 747–754, doi:10.1038/s41586-023-05857-4 (2023).

5 Reyes, J. M. et al. Hematologic DNMT3A reduction and high-fat diet synergize to promote weight gain and tissue inflammation. iScience 27, 109122, doi:10.1016/j.isci.2024.109122 (2024).

6 Pasupuleti, S. K. et al. Obesity-induced inflammation exacerbates clonal hematopoiesis. J Clin Invest 133, doi:10.1172/JCI163968 (2023).

7 Tobias, D. K. et al. Clonal Hematopoiesis of Indeterminate Potential (CHIP) and Incident Type 2 Diabetes Risk. Diabetes Care 46, 1978–1985, doi:10.2337/dc23-0805 (2023).

8 Bick, A. G. et al. Inherited causes of clonal haematopoiesis in 97,691 whole genomes. Nature 586, 763–768, doi:10.1038/s41586-020-2819-2 (2020).

9 Brunetti, L., Gundry, M. C. & Goodell, M. A. DNMT3A in Leukemia. Cold Spring Harb Perspect Med 7, doi:10.1101/cshperspect.a030320 (2017).

10 Venugopal, K., Feng, Y., Shabashvili, D. & Guryanova, O. A. Alterations to DNMT3A in Hematologic Malignancies. Cancer research 81, 254–263, doi:10.1158/0008-5472.CAN-20-3033 (2021).

11 Feng, Y. et al. Hematopoietic-specific heterozygous loss of Dnmt3a exacerbates colitis-associated colon cancer. J Exp Med 220, doi:10.1084/jem.20230011 (2023).

12 Guryanova, O. A. et al. Dnmt3a regulates myeloproliferation and liver-specific expansion of hematopoietic stem and progenitor cells. Leukemia 30, 1133–1142, doi:10.1038/leu.2015.358 (2016).

13 Guryanova, O. A. et al. DNMT3A mutations promote anthracycline resistance in acute myeloid leukemia via impaired nucleosome remodeling. Nature medicine 22, 1488–1495, doi:10.1038/nm.4210 (2016).

14 Jakobsen, N. A. et al. Selective advantage of mutant stem cells in human clonal hematopoiesis is associated with attenuated response to inflammation and aging. Cell Stem Cell 31, doi:10.1016/j.stem.2024.05.010 (2024).

15 Xie, X. et al. Clonal hematopoiesis and bone marrow inflammation. Transl Res 255, 159–170, doi:10.1016/j.trsl.2022.11.004 (2023).

16 Chalasani, N. et al. The diagnosis and management of nonalcoholic fatty liver disease: Practice guidance from the American Association for the Study of Liver Diseases. Hepatology 67, 328–357, doi:10.1002/hep.29367 (2018).

