## Supplementary Information for "Clonal hematopoiesis driven by *Dnmt3a* mutations promotes metabolic disease development"

**Supplementary Figure 1**
Dnmt3a Deficiency or Mutation in Hematological Cells Promotes Obesity and Diabetes

**Supplementary Figure 2**
Dnmt3a Deficiency or Mutation in Hematological Cells is Associated with inflammation and myeloid cell infiltration

**Supplementary Methods**

**Supplementary Table 1**
Histological grading criteria for steatohepatitis in mice

**Supplementary Table 2**
List of antibodies used in this study

**Supplementary References**

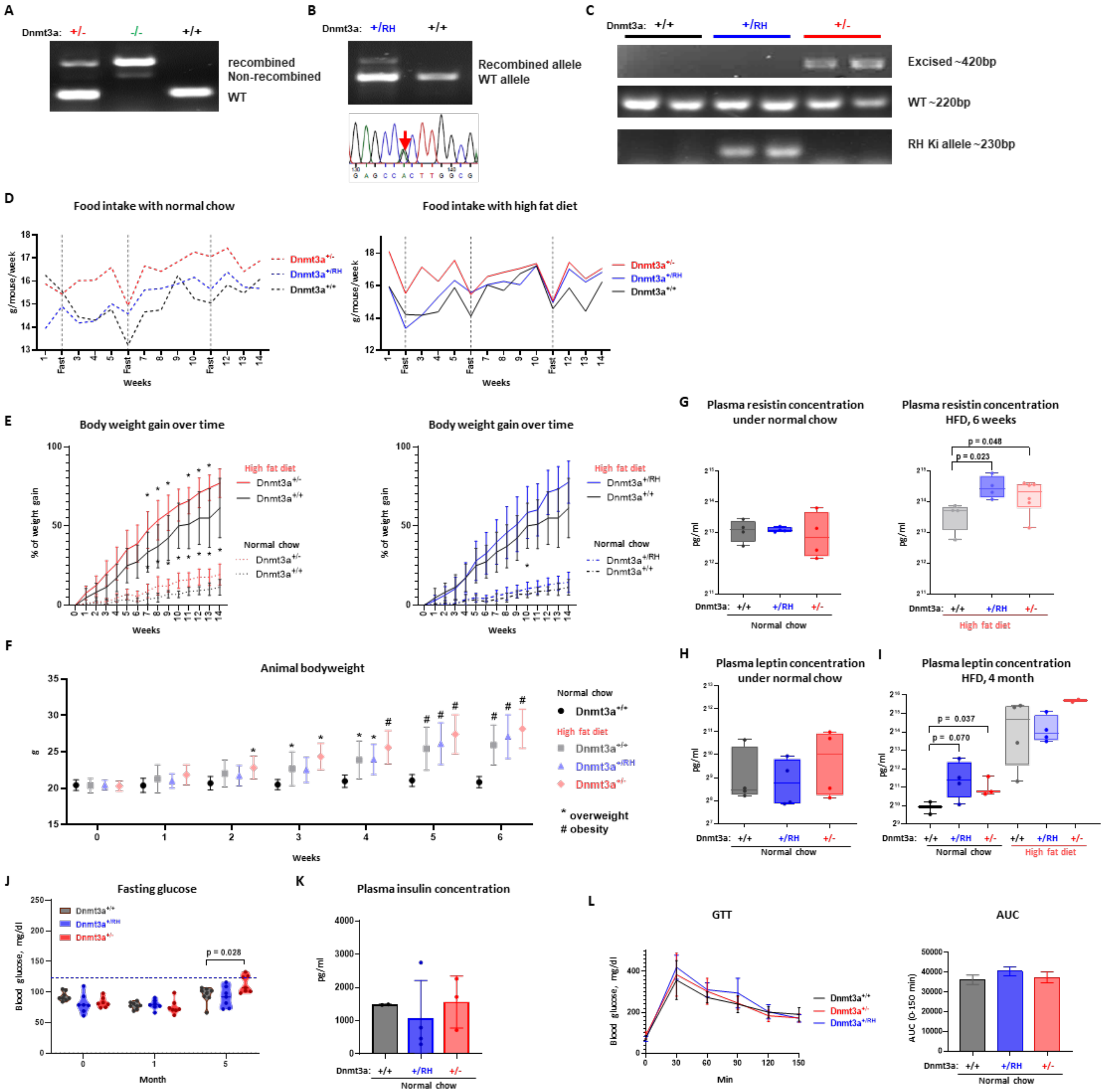


**Supplementary Figure 1. Hematopoietic-specific *Dnmt3a* alterations in a model of CH promote obesity and impaired glucose tolerance**

(A-B) Genotyping of donor mice confirms full recombination of the inducible knock-out or knock-in alleles. *Dnmt3a*$^{+/-}$ was genotyped using PCR of genomic DNA purified from blood nucleated cells (A). *Dnmt3a*$^{+/RH}$ was genotyped using both genomic DNA PCR and Sanger sequencing of peripheral blood nucleated cell cDNA (B). (C) Genotyping of peripheral blood from recipient mice 4 month after BMT confirms consistent presence of donor-derived cells. (D) Animal food intake, averaged per cage housing 5 mice of the same experimental group. (E) Animal body weight gain over time. (F) Animal body weight (n≥7). Mice are considered as overweight when their body weight exceeds that of age- and sex-matched controls by 10–20% an obese when it is more than 20%. (G) Plasma resistin levels in indicated groups. Plasma samples were collected at 14 weeks post BMT (6 weeks after diet randomization). (H-I) Plasma leptin levels in indicated groups, 6 weeks (H) or 4 months (I) after randomization to HFD or control chow. (J) Fasting blood glucose levels at baseline and at 1 or 5 months after diet randomization in a control normal chow group. (K) Plasma insulin concentration at 4 months after diet randomization in control normal chow group. (L) Glucose tolerance test (GTT) and calculated area under the curve (AUC) in indicated groups under normal chow; n≥5 for all groups.

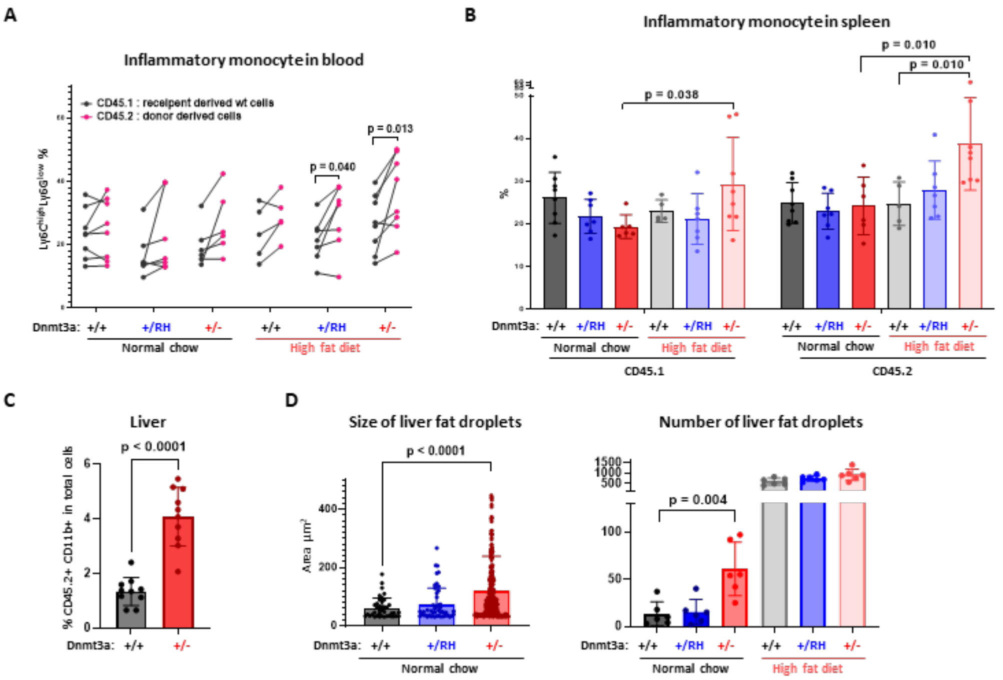


**Supplementary Figure 2. Model CH with *Dnmt3a* deficiency or mutation promotes inflammation, advanced steatohepatitis and liver damage**

(A-B) Proportion of inflammatory monocytes in CD45.1$^+$CD11b$^+$ or CD45.2$^+$CD11b$^+$ population in peripheral blood, paired Student's *t*-test (A) and in spleens, unpaired Student's *t*-test with Welch's correction (B). (C) Percentage of CD45.2$^+$ myeloid cells in liver, Student's *t*-test. (D) Size and number of liver fat droplets (n=3 per group, 2 fields per animal, Mann-Whitney rank sum test (left) and unpaired Student's *t*-test (right)).

### Methods

#### Animals

All animal experiments were conducted in accordance with ethical guidelines and were approved by the Institutional Animal Care and Use Committee (IACUC) at the University of Florida (UF). Mice were housed in a specific pathogen-free animal care facility with a controlled housing temperature (23°C) and 12-hour light/12-hour dark cycle. Conditional *Dnmt3a* knockout (*Dnmt3a*$^{+/fl}$), *Dnmt3a*$^{+/R878H}$ (corresponds to the *DNMT3A* R882H mutation in humans) knock-in, and *Dnmt3a*$^{+/+}$ mouse lines with *Mx1-Cre* deleter on a C57BL/6J background were generated via in-house breeding[1-3]. To achieve inducible hematopoietic-specific excision, 8- to 12-week-old *Dnmt3a*$^{+/R878H}$:*Mx1-Cre* and *Dnmt3a*$^{+/fl}$:*Mx1-Cre* mice received five intraperitoneal injections of poly(I:C) (InvivoGen, 20mg/kg of body weight). Poly(I:C)-treated *Dnmt3a*$^{+/+}$:*Mx1-Cre* mice are used as controls. *Mx1-Cre*–driven recombination was validated by PCR using genomic DNA purified from peripheral blood mononuclear cells two weeks after the final injection. Successful expression of the *Dnmt3a*$^{R878H}$ point mutant is further verified by Sanger sequencing of the peripheral blood mononuclear cell cDNA. All recipient mice used for bone marrow transplantation in this study were C57BL/6.SJL (CD45.1) immune-competent, healthy females procured from the Jackson Laboratory. Animals had *ad libitum* access to water and either high-fat (42 kcal% fat, TD.88137, Envigo) or standard chow (Teklad 2918 18% protein, Envigo) rodent diet.

#### Bone marrow transplantation (BMT)

Bone marrow (BM) cells were isolated from C57BL/6 (CD45.2) age-matched, fully recombined *Dnmt3a*$^{+/fl}$, *Dnmt3a*$^{+/R878H}$, and *Dnmt3a*$^{+/+}$ control donors, as well as from congenic C57BL/6.SJL (CD45.1) mice as wild-type support BM. As indicated in Figure 1A, lethally irradiated (5.4 Gy, twice) 6-week-old CD45.1 recipient mice received 20% individual genotype donor BM cells (CD45.2) and 80% wild-type CD45.1 BM cells through a tail vein injection ($2\times10^5$ CD45.2$^+$ donor BM cells and $8\times10^5$ wild-type CD45.1$^+$ BM cells with a total of $1\times10^6$ BM cells/recipient mouse). Animals were maintained on a normal chow diet for 8 weeks to allow full engraftment. After hematopoietic reconstitution was confirmed by peripheral blood flow cytometry analysis for CD45.1/CD45.2 chimerism, major mature lineages, and by CBCs at 8 weeks post-BMT, animals were randomly assigned to high fat diet or remained on normal chow.

**Flow cytometry analysis**

The blood and spleen single-cell suspensions were lysed with an RBC lysis buffer and then resuspended in phosphate buffered saline (PBS) containing 0.2% bovine serum albumin (BSA) (Sigma). The cells were then stained with fluorochrome-conjugated antibodies and analyzed by multiparameter flow cytometry performed using LSR Fortessa instrument (BD Biosciences). Data were analyzed using FlowJo software (v10.4.1). Antibodies used in this study are listed in Supplementary Table 2.

**Plasma cytokine and metabolic hormone analysis**

Plasma was separated from peripheral blood and sent to Eve Technologies (Canada) for mouse cytokine/chemokine and metabolic hormone (array MD44-plex and MRDMET12) analysis. Results are expressed as pg/mL of plasma.

**Glucose Tolerance Test**

Animals were fasted overnight, and glucose was administered by intraperitoneal injection at a dose of 1.5 g/kg. Blood glucose levels were measured using a blood glucose meter (AgaMatrix) immediately prior to dosing (0 min) and at 15, 30, 60, 90, 120, and 150 minutes post-loading via tail vein bleeding. Data are presented as mean ± standard error. Statistical difference between group means compared to *Dnmt3a*$^{+/+}$ control group at individual time point was assessed by Student's *t*-test; with $p<0.05$ considered significant ($n \geqslant 6$ animals per group).

**Histological analysis by H&E and Masson's trichrome staining**

Animals were euthanized by $CO_2$ asphyxiation, and harvested tissues were immediately fixed in 10% neutral buffered formalin solution (Sigma, MFCD00003274) for 24 hours. The samples were then submitted to the UF Molecular Pathology Core for processing. Tissues were paraffin-embedded, sectioned, and stained following standard protocols. Slides were imaged using Olympus VS200 Slide Scanner. White adipocyte size was quantified by measuring the cell area using "Analyze Particles" function in ImageJ software (v1.53k). Cells touching the image borders were excluded from the analysis to ensure accurate measurements. Steatohepatitis was evaluated through histological grading according to modified CRN criteria[4,5] (Supplementary Table 1). Liver fibrosis was assessed by quantifying blue-stained collagen positivity of Masson's trichrome staining by measuring area fraction of blue channel with ImageJ software (v1.53k). Statistical analysis was performed using two-tailed unpaired Student's *t*-test.

**Statistical analysis**

Animal bodyweight gain was analyzed using mixed-effects analysis. Unpaired comparisons between two groups were performed using Student's *t*-test for normally distributed data with equal variances, Welch's correction for unequal variances, or the Mann–Whitney *U*-test or Kolmogorov-Smirnov test for non-normal data or unequal variances, as appropriate. Data normality was tested using the Shapiro-Wilk test. Pairwise comparisons within the same mouse, comparing the inflammatory monocyte percentage in the CD45.1$^+$CD11b$^+$ and CD45.2$^+$CD11b$^+$ populations in the spleen and peripheral blood, were conducted using paired Student's *t*-test. Area under curve (AUC) was calculated using the trapezoidal rule. The lowest glucose value for each mouse was subtracted from the AUC. All statistical analyses were carried out with GraphPad Prism 10.3.1.

**Supplementary Table 1. Histological grading criteria for steatohepatitis in mice**

| | |
|---|---|
| ***Steatosis*** | |
| Grade 0 | <5% of hepatocytes |
| Grade 1 | < 33% of hepatocytes |
| Grade 2 | 33-66% of hepatocytes |
| Grade 3 | > 66% of hepatocytes |
| ***Inflammation*** | |

| | |
|---|---|
| Grade 0 | Absent |
| Grade 1 | <100 inflammatory cells per focus or <3 inflammatory foci per 20× field |
| Grade 2 | 100-500 inflammatory cells per focus or 3-4 inflammatory foci per 20× field |
| Grade 3 | >500 inflammatory cells per focus or >4 inflammatory foci per 20× field |
| ***Hepatocyte Injury/Ballooning*** | |
| Grade 0 | Absent |
| Grade 1 | Rare balloon cells |
| Grade 2 | Widespread hepatocyte ballooning and apoptosis |
| ***Modified NASH activity score (NAS)*** | |
| Grade 1 | 0-2 points |
| Grade 2 | 3-4 points |
| Grade 3 | 5-6 points |
| Grade 4 | 7-8 points |

Modified from Wong. *et al.*, which was based on NASH Clinical Research Network scoring system for NAFLD[4,5].

**Supplementary Table 2. List of antibodies used in this study.**

| Antibody | Fluorophore | Company | Catalog number | Dilution |
|---|---|---|---|---|
| CD45.1 | PacBlue | BioLegend | 110722 | 1:200 |
| CD45.2 | BV605 | BioLegend | 109841 | 1:200 |
| B220 | PE-Cy7 | BioLegend | 103222 | 1:200 |
| CD11b | AF700 | BioLegend | 101222 | 1:200 |
| Gr1 | APC | BioLegend | 108412 | 1:200 |
| Ly6G | BV785 | BioLegend | 127645 | 1:200 |

**References**


1. Guryanova, O. A. *et al.* DNMT3A mutations promote anthracycline resistance in acute myeloid leukemia via impaired nucleosome remodeling. *Nat Med* **22**, 1488-1495, doi:10.1038/nm.4210 (2016).
2. Venugopal, K. *et al.* DNMT3A Harboring Leukemia-Associated Mutations Directs Sensitivity to DNA Damage at Replication Forks. *Clin Cancer Res* **28**, 756-769, doi:10.1158/1078-0432.CCR-21-2863 (2022).
3. Feng, Y. *et al.* Hematopoietic-specific heterozygous loss of Dnmt3a exacerbates colitis-associated colon cancer. *J Exp Med* **220**, doi:10.1084/jem.20230011 (2023).
4. Chalasani, N. *et al.* The diagnosis and management of nonalcoholic fatty liver disease: Practice guidance from the American Association for the Study of Liver Diseases. *Hepatology* **67**, 328-357, doi:10.1002/hep.29367 (2018).
5. Wong, W. J. *et al.* Clonal haematopoiesis and risk of chronic liver disease. *Nature* **616**, 747-754, doi:10.1038/s41586-023-05857-4 (2023).
